# Democratizing Organ-on-Chip Technologies with a Modular, Reusable, and Perfusion-Ready Microphysiological System

**DOI:** 10.1101/2025.04.30.651503

**Authors:** Daniel J. Minahan, Katherine M. Nelson, Filipa Ribeiro, Bryan J. Ferrick, Alexandra M. Zurzolo, Kira Byers, Jason P. Gleghorn

**Affiliations:** Department of Biomedical Engineering, University of Delaware, Newark, DE 19716; Department of Chemical and Biomolecular Engineering, University of Delaware, Newark, DE 19716; Department of Mechanical Engineering, University of Delaware, Newark, DE 19716

**Author notes:** Authors contributed equally (Daniel J. Minahan, Katherine M. Nelson).

**Keywords:** Modular Microfluidics, Organ-on-Chip (OOC), Microphysiological System (MPS), *In Vitro* Tissue Models, Low-Resource

## Abstract

Organ-on-chip (OOC) technologies, also called microphysiological systems (MPS), offer dynamic microenvironments that improve upon static culture systems, yet widespread adoption has been hindered by fabrication complexity, reliance on poly-dimethylsiloxane (PDMS), and limited modularity. Here, we present a modular MPS platform designed for ease of use, reproducibility, and broad applicability. The system comprises layered elastomeric inserts for dual monolayer cell culture, which is clamped within a reusable acrylic cassette for perfusion studies. This enables researchers to decouple model establishment from flow experiments and streamline their workflows. We validated the system using dual epithelial and endothelial cell co-culture under static and perfused conditions, including shear-induced alignment of HUVECs. Material testing confirmed biocompatibility, while vinyl cutting reproducibility demonstrated high manufacturing fidelity. The platform reliably supported long-term culture (up to 14 days), and the open insert format facilitated uniform seeding and imaging access. This approach enables parallelized experimentation, minimizes pump usage, and is well-suited for labs without microfabrication infrastructure. By combining fabrication flexibility with biological robustness, this work establishes a generalizable platform for modular tissue-chip development adapted to diverse organ systems and serves as a foundational framework for democratizing advanced *in vitro* model systems.

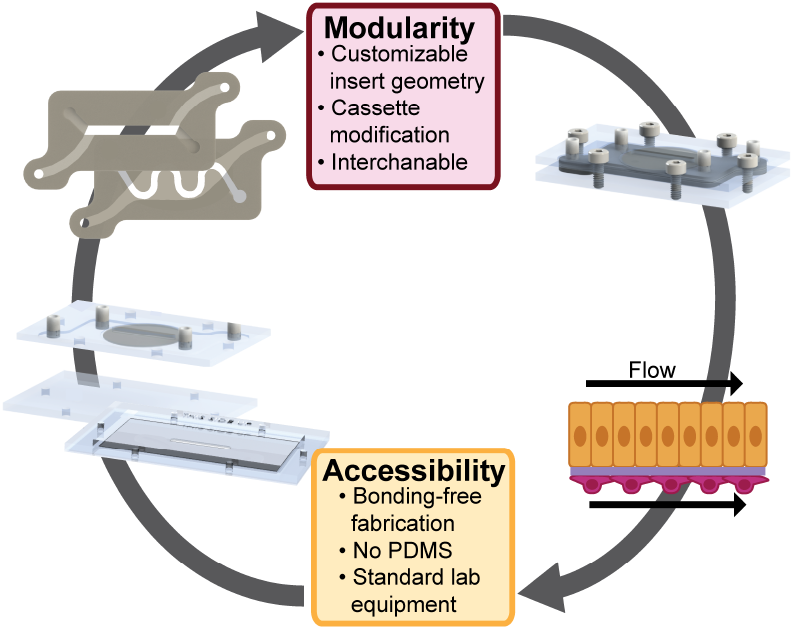

## Introduction

Microphysiological systems (MPS) have emerged as valuable tools for *in vitro* modeling of tissue physiology and disease by incorporating flow, shear stress, and multicellular coculture in microscale environments (1–4). However, many existing organ-on-chip (OOC) devices rely on polydimethylsiloxane (PDMS) and soft lithography (3, 5, 6), requiring access to cleanroom infrastructure and technical expertise that remains out of reach for many biological laboratories. Rapid prototyping techniques such as 3D printing, laser cutting, and other smart craft cutting methods have expanded manufacturing accessibility due to their relatively low cost and ease of use. These methods can result in fast manufacturing and precise geometries at low cost and have been used to create MPS systems (7–14). However, like other fabrication methods, these systems are irreversibly sealed, which limits the types and number of assays that can be run with one device. These constraints limit widespread adoption and hinder integration into non-engineering-focused biological labs.

To address this barrier, we developed a modular, PDMS-free MPS platform fabricated using benchtop laser and vinyl cutting tools, with no bonding or cleanroom steps. The platform consists of a reusable acrylic ‘cassette’ and compressible elastomeric ‘inserts’ that can be easily interchanged or customized for different experimental designs and seal fluidic channels upon clamping. This compression reversibly forms leak-tight fluid channels without the need for adhesives or thermal bonding. The reusable cassette can be fabricated to include glass coverslips, which create a system that can be live-imaged via microscopy and can also be sterilized for multiple uses. The consumable insert was fabricated with a commercially available track-etched porous membrane and acts simultaneously as the cell culture scaffold and the gasket, simplifying assembly while enabling reversible use and customizable geometry. The system is designed to replicate key features of commonly used Transwell inserts - enabling ease of seeding, straightforward cell culture workflows, and compatibility with standard biological lab tools - while offering the benefits of dynamic microfluidic perfusion. By decoupling model formation from perfusion, users can seed multiple inserts in parallel and initiate flow only when needed. By reducing the technical and financial barriers to entry, this platform empowers broader adoption of dynamic *in vitro* models for mechanistic studies and facilitates the iterative workflows required for translational discovery.

## Materials and Methods

### Device fabrication

**Insert:** For silicone inserts, double-sided adhesive film (3M, 9500PC) was placed on one side of 1/32” silicone sheets (McMaster-Carr, 86465K21), taking care not to create air pockets between the layers. Any large air bubbles present were gently rolled out using a western blot roller (SNAP2RL, Millipore-Sigma). With the paper backing on, the laminated silicone and adhesive were fed into a vinyl cutter (USCutter, SC or Titan3), silicone side up. The upper and lower insert patterns (made in SolidWorks, converted to .SVG in Adobe Illustrator) were traced with a 45° blade at 100 mm/s and 300 g of pressure (SC) or 12 mm/s and 280 g of pressure (Titan 3). Inserts were assembled by sandwiching a 3 µm pore size polycarbonate membrane (Cytiva, 10418306) between two cut silicone sheets (**Figure 1A**). Inserts were sterilized using UV Ozone (BioForce Nanosciences, ProCleaner). Inserts were placed within the sterilization chamber and exposed to ozone and ultraviolet (UV) light for 5 minutes per side to ensure thorough surface sterilization. The inserts were then either transferred to a sterile container for immediate use or sealed in a sterile container for storage until future use.

**Fig. 1.**
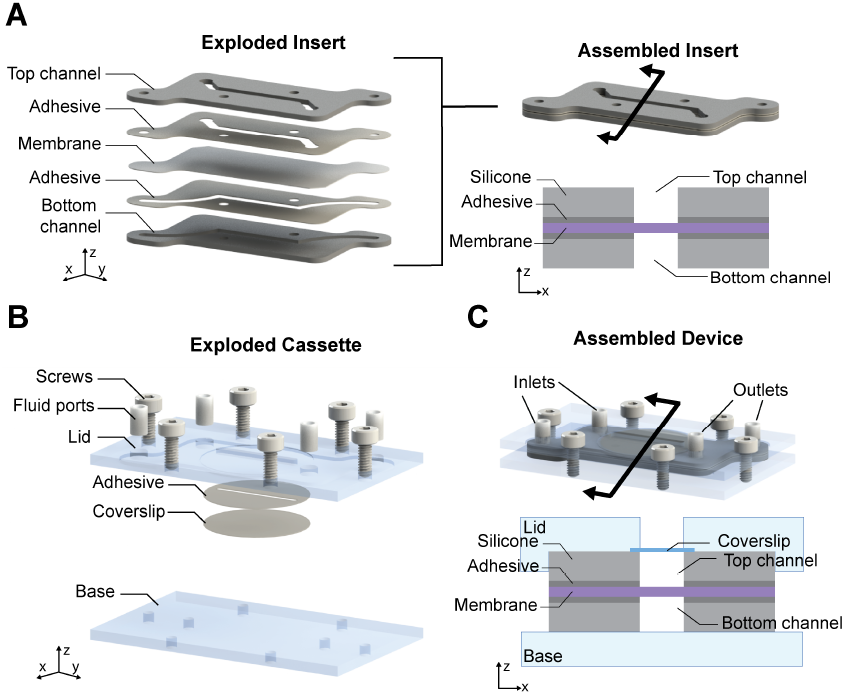
Overview of modular MPS platform. **(A)** The insert is made from stacked silicone pieces, a track-etched membrane, and double-sided adhesive film for cell culture. **(B)** The cassette is comprised of a top and bottom acrylic piece with inset coverglass for imaging. **(C)** The insert is clamped into the cassette using screws to create fluidic channels and fluid ports to connect external pumps.

**Cassette:** The cassette has two parts – a lid and a base both made from acrylic sheets of various thicknesses (McMaster-Carr) laser cut using a 40W CO2 laser and a Laser Muse 1.5” ZnSe Focus Lens (Full Spectrum Laser, Hobby Series 20x12). The standard cassette allowed for imaging via an upright microscope, while the inverted imaging cassette allowed for high magnification imaging with an inverted microscope. <u>Standard Cassette:</u> The lid was made from a 3/16” acrylic sheet (McMaster-Carr, 8560K211). The acrylic was rastered twice for an insert recess and a circular glass coverslip (No 1.5, Ø15 mm). Vector cut clearance holes allowed for M2 screw placement. In the raster layer containing the glass coverslip, the area was sanded flat or smoothed with heat to prevent coverslip breakage, and a vinyl cut adhesive sheet (3M 9500PC) was adhered. The circular coverslip was then adhered to the adhesive. Tubing was cut and glued to the inlet and outlet ports to connect to hypodermic blunt tips and the pump. For the standard cassette base, vector cut holes were made in a 3/16” acrylic sheet (McMaster-Carr, 8560K211) to allow tapping of M2 screws (**Figure 1B, 2A**). <u>Inverted Imaging Cassette:</u> For the inverted imaging cassette lid, vector cut holes were made in a 3/16” acrylic sheet (McMaster-Carr, 8560K211) to allow tapping of M2 screws and tubing holes, and a raster cut for an insert recess. Tubing was cut and glued to the inlet and outlet ports to connect to hypodermic blunt tips and the pump. The imaging cassette base was comprised of three acrylic sheets welded together with a coverslip. Two 0.01” acrylic films (McMaster-Carr, 4076N11) were vector cut for screw holes and central viewing ports. A third 1/16” acylic piece (McMaster-Carr, 8560K171) was vector cut to shape with screw holes. The cassette base parts were then assembled via acrylic weld by stacking the thicker piece on top of the two identical pieces. A vinyl cut adhesive sheet (3M 9500PC) was adhered to the resulting inset well that was created, and a rectangular coverslip (McMaster-Carr, 1149T13) was adhered to the adhesive, creating a cassette base that allows for *in situ* imaging on an inverted microscope (**Figure 2B**).

**Fig. 2.**
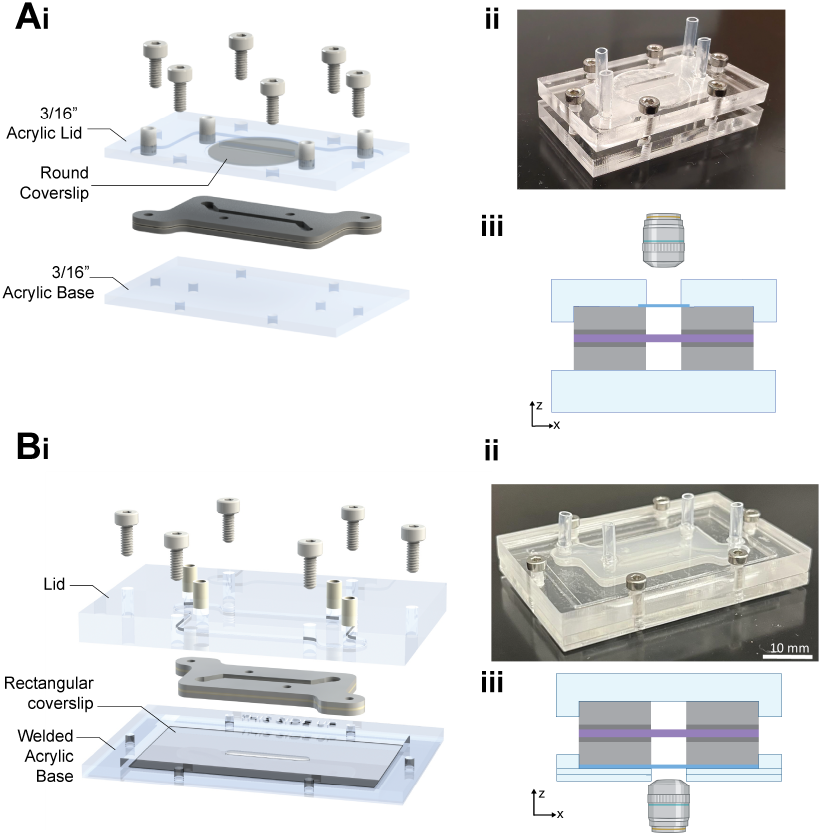
The cassette is modular and can be adapted for *in situ* imaging in upright or inverted microscopy configurations. **(A)** The standard cassette has a thick acrylic base and a coverslip in the lid, enabling upright microscopy. **(B)** The inverted imaging cassette is designed to accept the same insert, but optimized for higher magnification/resolution imaging on an inverted epifluorescent or confocal microscope. The acrylic base is thinner, houses the coverslip, and has greater access to the objective to decrease the working distance needed for *in situ* or timelapse imaging. “THIS SIDE UP” is etched into the base to aid in user orientation during model assembly. **i:** CAD rendering, **ii:** photograph of assembled devices, **iii:** cross-section schematic and imaging orientation.

The inserts could be reversibly clamped into the cassette using screws. Food dye was used in the resulting microfluidic channels to visualize channels and proper sealing (**Figure 1C, 2, 4AB**).

### Vinyl cutting reproducibility

Surface topography, height, and width measurements of the insert were obtained using a Keyence VK-X3000 Surface Profiler. At least three locations in each channel were imaged from at least four channels cut from three different batches of silicone. Each batch of silicone was cut by a different person at a different time. Samples were imaged using a 5X objective lens with both laser and optical imaging modes to capture surface features and to generate high-resolution 3D surface profiles. The height was measured from the bottom-most location to the edge of the channel. The width was measured at the bottom of the channel. Height and width measurements were extracted from the 3D profile data using the VK-X 3000 MultiFileAnalyzer software (Keyence). Measurements were compared to determine intra-batch and between-batch variability, as well as compared against CAD designs.

### Material toxicity assessment

Strips (40 mm x 12 mm) of PDMS (control), silicone sheet, and the double-sided adhesive film were incubated in 10mL of F-12K media (ATCC) at 37°C for 10 days. BeWo b30 placental cells were plated at 20,000 cells/well in 96-well plates and cultured overnight in conditioned medium. The following day, alamarBlue was added to the media and incubated for 3 hours. The fluorescence (560/590 ex/em) of each well was then read using a plate reader (BioTek Synergy H1M) to establish a baseline for each condition. After measurement, conditioned medium was added to the wells, and AlamarBlue was used to measure metabolic activity at 6, 19, and 26 hours. The fluorescence at each time point was plotted and normalized to an unconditioned (without any materials incubated) media control.

### Cell culture in insert and characterization

#### Cell seeding

Human umbilical vein endothelial cells (HUVECs, Lonza) were cultured in endothelial cell growth media (EGM-2, Lonza) and used between passages P2-P10. Madin-Darby Canine Kidney cells (MDCKs, ATCC) were cultured in Dulbecco’s Modification of Eagle’s Medium (DMEM, Corning) and used between passages P30-P40.

A PDMS well was created to place the insert on during cell culture (**Figure 3**). The polycarbonate membrane of the insert was coated with 0.1% gelatin for 45 minutes at 37°C on both sides, after which the excess gelatin was removed. HUVECs were seeded (150,000 cells/insert) on the upward-facing channel by pipetting and allowed to adhere for 4 hours. After 4 hours, EGM-2 media was added to the PDMS well and the HUVECs were cultured for 24 hours. The insert was then flipped, and MDCK cells were seeded (160,000 cells/insert) on the now empty upward-facing channel. Inserts were statically cultured until HUVEC and MDCK monolayers formed. After 8 days, monolayer formation was confirmed by E-Cadherin, VE-Cadherin, and phalloidin immunofluorescent staining.

**Fig. 3.**
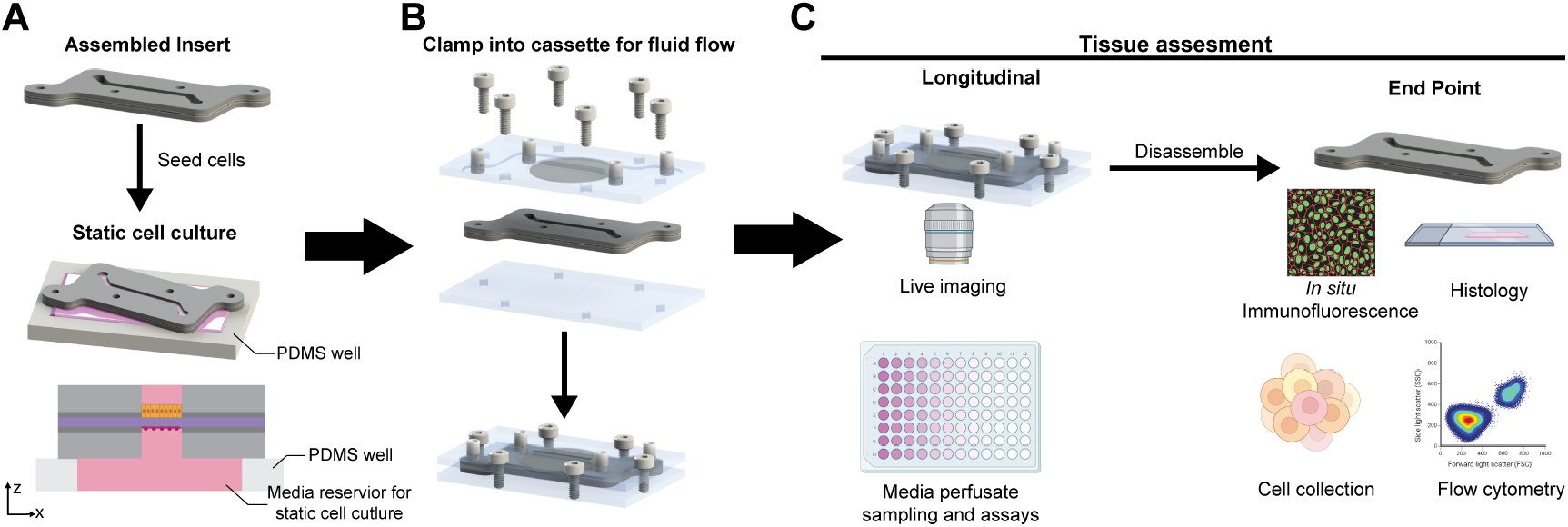
Workflow of cell culture, perfusion, and experimental assays in the device. **(A)** The insert is seeded with the desired cells onto the membrane using a pipette on a PDMS well. Media fills the well to create a larger media reservoir under the insert. After 24 hours of culture, the insert can be flipped, and a second cell type can be seeded onto the membrane. Inserts are cultured statically for 2-4 days to establish cell monolayers. This procedure and static cell culture assembly is similar to those used for a standard Transwell cell culture insert. **(B)** After the tissue model is established, the insert is clamped into the cassette, and the system can be connected to a pump to establish culture with relevant fluid flows for desired experimental conditions. **(C)** Longitudinal assays can be completed in situ, including timelapse imaging and perfusate analysis, or the device can be disassembled, and standard endpoint assays can be completed.

As a comparison, standard Transwell cell culture inserts were coated with 0.1% gelatin for 45 minutes at 37°C, after which the excess gelatin was removed. HUVECs or MD-CKs were seeded on the insert surface of the apical compartment at a density of 100,000 cells/cm^2^. EGM-2 (HUVECs) or DMEM (MDCKs) media was added to the basal compartment well. Transwells were cultured until HUVEC and MDCK monolayers formed, around 7-8 days, as confirmed by E-Cadherin, VE-Cadherin, and phalloidin immunofluo-rescent staining (15).

#### Perfusion

Inserts were clamped into cassettes after static culture and connected to a peristaltic pump. Flow was initiated at 1.27 uL/min (1.36x10^-3^ dyne/cm^2^) for the MDCK channel and 30 uL/min (0.03 dyne/cm^2^) in the HUVEC channel. Flow was progressively ramped to 3 mL/min (3.2 dyne/cm^2^) over the span of 48 hours for the HUVEC channel to induce a low physiological shear stress and HUVEC alignment. Perfusion was maintained over 14 days of culture. For experiments comparing static and flow conditions, inserts were cultured in a 10 cm petri dish statically in parallel with inserts in the cassette under flow.

#### Immunofluorescent staining

Following culture, inserts were removed from the cassette and cells were fixed with 4% paraformaldehyde for 1 hour at room temperature, permeabilized with 0.1% Triton-X for 2 hours at room temperature, and blocked overnight at 4°C in blocking buffer (0.2% gelatin, 0.5% BSA, 0.1% Tween-20) (6, 16). The actin cytoskeleton and nuclei were directly stained via phalloidin and Hoechst 33342 (Invitrogen), respectively, while VE-cadherin (Santa Cruz Technology) and E-cadherin (Santa Cruz Technology) were visualized via antibodies. Fluorescent images of immunofluorescently stained cells were captured using a Zeiss LSM 800 confocal microscope.

#### HUVEC alignment analysis

As a functional assay, inserts were cultured for 48 hours under flow and fixed and stained. Static cultures were kept outside of the cassette in a 10 cm petri dish for 48 hours to match the flow conditions. Alignment was quantified from immunostained HUVECs using ImageJ Ellipse Fitting for 3 fields of view. Cells were considered aligned if the angle between the cell axis and channel wall was <45° or >135°.

## Results

### Generation of a multilayer modular organ-on-a-chip

We designed and created a modular MPS, which consisted of reusable laser-cut acrylic cassettes (lid and base) and a layered compressible insert (**Figure 1, 2**). The insert and cassette platform enabled the formation of stacked fluidic channels separated by a synthetic track-etched membrane by clamping the compressible insert within the cassette secured by screws (**Figure 3**). Fluid ports within the lid connect tubing to a commercially available peristaltic pump, allowing for the independent perfusion of media through the top and bottom channels, as is common in other PDMS MPS systems. The standard cassette provided clamping force to seal the fluidic channel during flow, but due to the geometry with a coverglass located in the top cassette lid, it was limited for *in situ* and timelapse imaging experiments. Therefore, we also designed an inverted imaging cassette that fits the same inserts and could be used for high magnification live imaging *in situ* during perfusion (**Figure 2**). The insert and cassette are compatible with UV ozone or 70% ethanol solution sterilization. The cassette was specifically designed to be made with a bench-top laser cutter and readily available commercial materials (acrylic, screws, tubing connectors) to enable ease of manufacturing and translation to other experimental fields.

The insert was fabricated from silicone sheets cut on a vinyl cutter and supported dual monolayer co-culture of cells on either side of the track-etched membrane, similar to a transwell insert. Importantly, due to the format of the insert-cassette platform, the insert channel geometry can be rapidly changed depending on the experimental requirements (**Figure 4AB**); the time to go from a CAD design to a fabricated functional insert was less than one hour. Channel geometries were cut from a CAD design on the vinyl cutter, and the channel height was defined by the thickness of the commercially available silicone and the double-sided adhesive film. Quantitative measurements of the channel depth demonstrate minimal batch-to-batch variance in depth, and all measurements are within the manufacturer’s tolerance for their products (**Figure 4C**). The consistency of channel depth between batches of manufactured inserts is valuable as minimal variance in channel depth ensures that there is consistent shear stress applied to the cell monolayers between devices for a given flow rate setting on the pump.

**Fig. 4.**
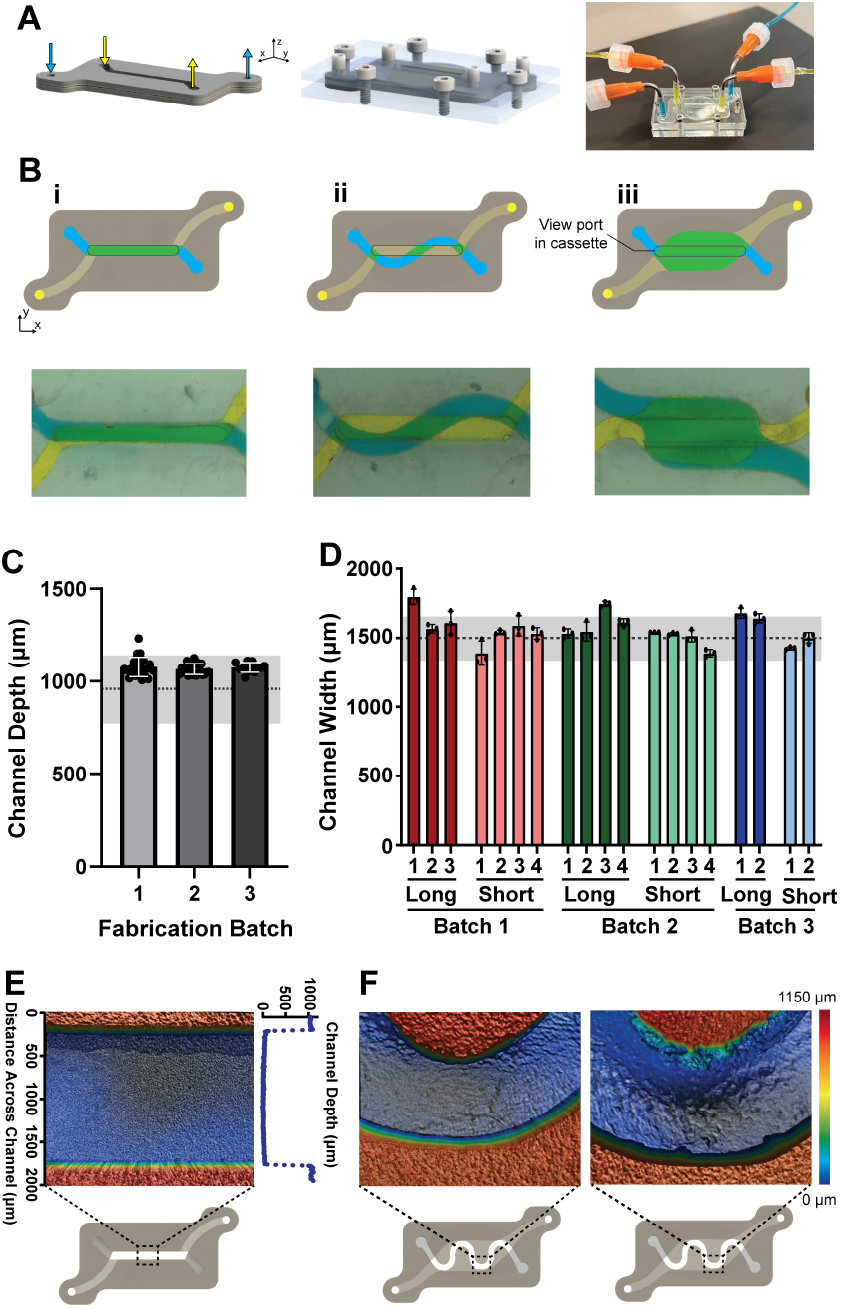
The modular insert can be rapidly made/prototyped using a benchtop vinyl cutter. **(A)** The insert supports the membrane for cell culture and defines the fluid channels when clamped into the cassette and connected to a pump. **(B)** Insert fluidic channels are rapidly prototyped and can be easily modified to fit within the boundary of the insert geometry to be universally compatible with the cassette design. Examples include a standard straight two-channel geometry **(i)**, one serpentine channel over a straight channel **(ii)**, and a differentially perfusable large overlapping cell area **(iii)**. Yellow and blue dyed water was perfused into the two channels of the devices to visualize each channel and their overlap (green). The microscopy view port in the cassette can also be seen. **(C)** The silicone sheet and double-sided adhesive tape thickness define the height of the channels (934*±* 178 *µ*m). There is limited variability in channel depth between channels in the same insert, between inserts, or between different fabricated batches of inserts compared to the expected channel height. n=24 for batch 1 and 2, n=12 for batch 3. The gray dotted line indicates the expected height of the channel based on the silicone and adhesive manufacturing heights, with the shaded area representing the manufacturer’s reported error on silicone sheet thickness. A nested one-way ANOVA indicated no significant difference between channel heights, *α* = 0.05. **(D)** The channel width of multiple inserts made in different batches. The gray dashed line (1500 *µ*m) represents the width from the CAD design, and the gray region is 10% difference. **(E)** Surface profilometry image of channels within fabricated inserts in the standard straight channel geometry with a corresponding plot of the channel depth. **(F)** Surface profilometry images of a curved channel geometry show that while the shape can be cut, error increases with channels of high radius of curvature with occasional cutting imperfections of the side-wall.

Given that the vinyl cutter was cutting a deformable silicone sheet, we also assessed differences in channel widths within the same batch cut from a single silicone sheet and between batches, and compared those to the intended dimensions from our CAD design (**Figure 4D**). We measured the widths at the bottom of the channel, which was the bottom surface when cutting, as we anticipated the deformable nature of the silicon sheet would be the most pronounced at that location. Channel widths were measured in three positions along the length and averaged to determine the mean channel width of either the long or short straight channel that make up the channels in the insert. Overall, there was minimal variance in width along a given channel, and all channels widths were no more than 25% away from the CAD-defined width (1500 *µ*m) (range 88-124%). We found that the error was more pronounced in longer channels, presumably due to the deformable nature of the silicone sheet; however, overall, most channels (84%) were no more than 10% from the expected width, confirming reproducibility. The the average width of the longer channel was more likely to be larger than the defined channel width (33% were more than 10% larger than the expected width), compared to the shorter channels (0% of the channels were more than 10% different from the expected width). Visual inspection from different regions of the channel in multiple inserts indicated that the cut channels were uniform and consistent in the standard straight channel insert (**Figure 4E**). However, in a second geometry with tight curvature in a channel, the channel was often cut correctly, but, on some occasions the cut was not perfect, indicating that while many geometries are possible, care needs to be taken to inspect the cuts, especially when geometries with tight curvature are used (**Figure 4F**). These methods enable rapid prototyping of tissue configurations and scalable manufacturing of inserts within the same cassette format, features that are needed for mechanistic studies relative to a fixed geometry for screening applications.

### Materials are non-toxic and support cell culture

Cytotoxicity assays revealed no adverse effects of the insert materials on epithelial cell metabolism. Using an alamarBlue assay, the metabolic activity of BeWo b30 epithelial cells cultured in material-conditioned media was measured and compared to cells cultured in unconditioned medium (**Figure 5**). Placental trophoblast BeWo b30 cells were used as they have high sensitivity to lechants and environmental chemicals and are sensitive to metabolic disturbances from the extracellular environment (17–20). While PDMS has been reported to leach unpolymerized polymer into media if not cured and rinsed properly (21, 22), we did not see any toxicity that resulted in metabolic changes from PDMS. None of the insert materials (silicon or the double-sided adhesive film) showed significant disruption of metabolic activity after 26 hours, indicating that the commercially available materials in the MPS insert are biocompatible for cell culture.

**Fig. 5.**
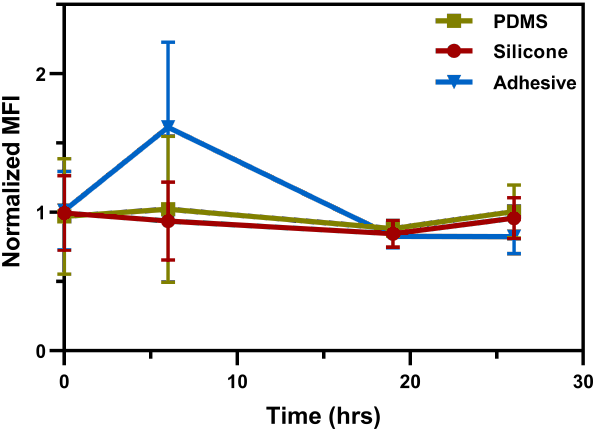
Toxicity study demonstrates biocompatibility of insert materials. alamarBlue assay measured the metabolic activity at t= 0, 6, 19, and 26 hours of incubation with conditioned media from silicone, PDMS, or the adhesive film for 10 days. Values were normalized to fresh medium. n=3. There was no statistical significance between materials at any time point based on a two-way ANOVA statistical test, *α* = .05.

### Cell culture is possible in static or dynamic conditions

We cultured HUVECs or MDCKs on a transwell cell culture insert (**Figure 6AB**) and compared their morphology to HUVECs and MDCK cells dual-cultured on either side of the MPS insert (**Figure 6CD**). Cells formed continuous monolayers and were morphologically similar on both systems, as evidenced with immunofluorescent staining of E-cad (MDCK) or VE-cad (HUVEC) along with nuclear staining using hoechst. In our MPS insert, co-cultured monolayers of MDCKs and HUVECs were visualized *in situ* on either side of the MPS insert membrane from reconstruction of z-stack images (**Figure 6E**), creating a multicellular tissue-level architecture consistent with other PDMS-based MPS devices (1, 23). Fluorescent imaging confirmed the monolayer’s integrity, and immunostaining demonstrated the presence of appropriate epithelial and endothelial markers for both models cultured under static conditions (24).

**Fig. 6.**
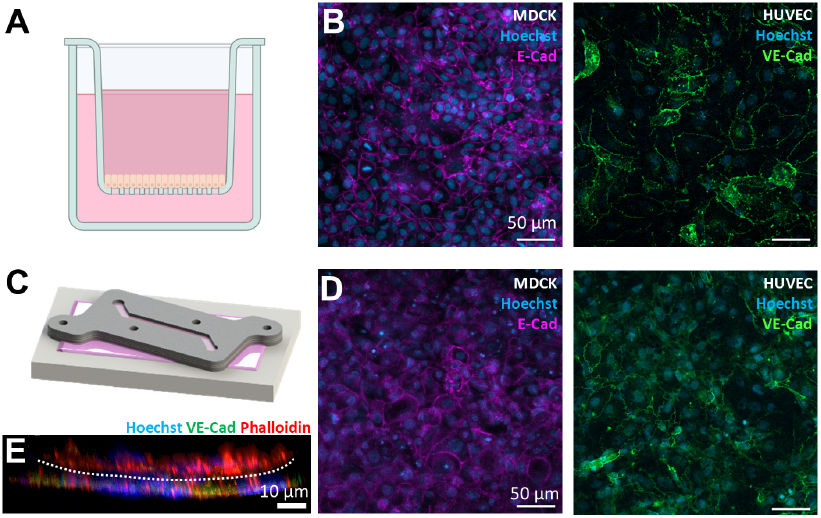
Comparison of static cell culture in a Transwell cell culture insert and the device insert over 8 days of culture show no phenotypic differences with intact cell monolayer formation. **(A)** MDCK epithelial and HUVEC endothelial cells were cultured statically in a standard Transwell insert and **(B)** immunofluorescently stained for nuclear and junctional markers. **(C)** Static culture using the device insert with corresponding **(D)** immunofluorescent staining of MDCKs and HUVECs on either side of an insert. **(E)** Z-stack reconstruction of the cross-section of the insert showing two cell layers on either side of the membrane (dotted line) in the insert.

After the establishment of the tissue model in the insert under static culture in a petri dish, the insert was clamped into the cassette to establish fluid flow and a relevant shear stress was applied to the epithelial and endothelial tissues via fluid flow (**Figure 7A**). To validate successful perfusion and appropriate phenotypic shifts were observed from shear stress, we perfused the HUVEC channel with media at 3.2 dyne/cm^2^ or the MDCK channel at 1.36x10^-3^ dyne/cm^2^ for 48 hrs. Both HUVECs and MDCKs were able to be stably cultured in the cassette under flow as seen by VE-cad and E-cad immunofluorescent staining, respectively (**Figure 7B**). We quantified HUVEC alignment of each endothelial cell within three fields of view in static and perfused conditions (**Figure 7C**), as HUVECs are known to align with the direction of flow (25). Approximately 70% of HUVECs aligned along the direction of flow (**Figure 7D**) in our modular MPS compared to only 40% of HUVECs which aligned along the channel length when cultured statically in the insert.

**Fig. 7.**
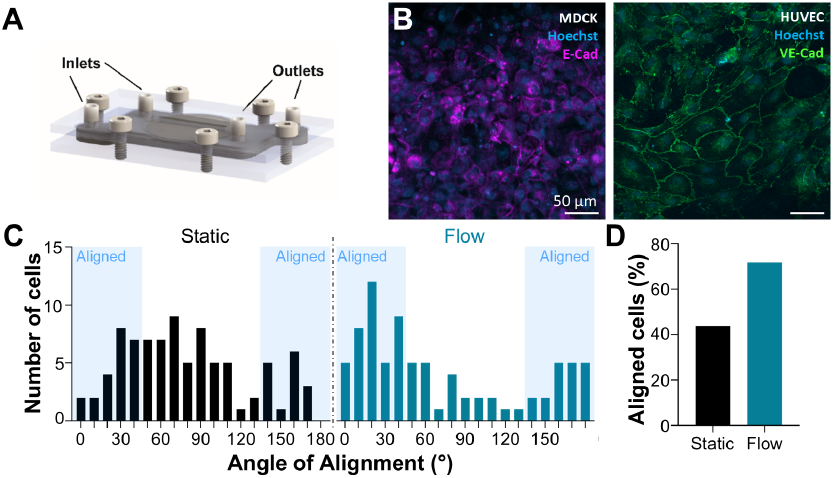
Cell phenotypic changes in HUVEC flow-mediated alignment occur within the MPS device when cultured in the cassette and perfused with medium. **(A)** MDCK epithelial or HUVEC endothelial cells were cultured under flow in the assembled device and **(B)** immunofluorescently stained afterward two days of culture in the cassette. **(C)** HUVEC alignment under static and flow culture conditions was quantified from three fields of view. **(D)** Flow-induced alignment of HUVECs validates a common phenotypic marker observed in traditional OOC devices.

The open format of the insert enabled straightforward and consistent cell seeding (∼100% success rate) and mimicked Transwell insert workflows, allowing multiple tissue models to be prepared in parallel. These inserts could then be clamped into the cassette when needed, reducing pump demand and supporting experimental staging. Devices were successfully cultured in perfusion for up to 14 days, with experimental termination based on study design rather than platform or tissue failure. Additionally, the ability to remove the insert from the cassette at the end of the experiment allowed for high-fidelity immunofluorescent imaging to enable the quantification of cell phenotype (**Figure 3**).

## Discussion

Microfluidic organ-on-chip (OOC) devices offer critical advantages over traditional static cultures by incorporating fluid flow and shear stress into cellular environments. However, the majority of existing platforms require advanced fabrication infrastructure, have material constraints that produce irreversibly bonded devices, and are difficult to modify without specialized expertise. These constraints have limited broader adoption, particularly in labs focused on biology, pharmacology, or translational research.

We present a modular, PDMS-free MPS platform designed to overcome these challenges through user-friendly fabrication, modular assembly, and flexibility in geometry and materials. Our system is reversibly clamped, eliminating the need for irreversible bonding and allowing for easier cell culture workflows, assay access, and endpoint imaging. The insert can be fabricated from commercial silicone and supports dual-cell monolayer culture on either side of a track-etched membrane, similar to a standard Transwell insert (12, 13). Devices were cultured successfully for up to 14 days, with termination based on experimental goals rather than platform failure, demonstrating the robustness of the system. As cell seeding occurs in the open insert format, successful monolayer formation is more likely than with fluid flow-based seeding methods. This significantly lowers the barrier to adoption and enhances reproducibility. Compared to soft lithography or commercial PDMS devices, this approach reduces costs, simplifies iteration, and accelerates model development.

The “insert” architecture decouples tissue model formation from flow application, allowing cells to be cultured under static conditions before introduction into the cassette for perfusion. This enables parallelized experiments, reduces pump demand, and increases experimental throughput, as perfusion assays can run concurrently with tissue model development. The ability to remove the cell culture platform from the fluidic system not only enables more applications but the reuse of the cassette also reduces costs. This decoupled format introduces a new degree of flexibility into organ-on-achip workflows, making longitudinal and multiplexed studies more tractable.

The modularity and tunability of our platform make it adaptable for diverse applications that have been demonstrated in other 3D MPS models (26–28). Channel width, length, geometry, and height can be customized to match experimental needs. We validated the reproducibility of vinyl cutting as a fabrication method for the insert. Across multiple batches, feature dimensions remained within tight tolerances relative to CAD specifications, supporting the feasibility of consistent multi-user or multi-site implementation. Compared to soft lithography, the fabrication of the cassette and insert avoids toxic reagents, cleanroom infrastructure, and expensive molds (29, 30). The system can be built using only a laser cutter, vinyl cutter, and standard lab tools, making it viable for widespread distribution or on-demand local production. Designing reusable components in a rapid-prototyped, easy-to-use system allows for the democratization of an MPS model, where multiple experiments can be run using a single base system. This biological modularity enables rapid iteration and configuration of tissue models, supporting single-cell type culture, dual monolayers, or more complex constructs without redesigning the entire device.

By prioritizing accessibility, customization, and reusability, our platform represents a framework for modular organon-chip development. It bridges a gap between engineered microenvironments and standard biological workflows, enabling researchers to build sophisticated tissue models without requiring specialized fabrication infrastructure. We anticipate that this platform will facilitate broader experimentation, iteration, and collaboration across domains — from academic biology labs to translational and therapeutic development.

## Conclusion

The advancement of microfluidic *in vitro* models has enabled precise control over dynamic environmental variables, offering improvements over traditional 2D culture for studying cell behavior and function. Yet, despite their scientific potential, many of these systems remain confined to expert bioengineering labs due to the complexity and cost of fabrication, operation, and modification. To address this barrier, we developed an accessible, modular MPS platform that is inexpensive, reversibly sealed, and fabricated from commercially available materials using standard laboratory tools. Its design enables widespread use, even in non-engineering settings, while maintaining compatibility with static and dynamic workflows, live imaging, and a range of biological applications. This platform can be readily adapted for use across diverse organ systems and research questions. By decoupling tissue model formation from perfusion and simplifying the fabrication process, it helps democratize organ-on-chip technologies, broadening access to advanced *in vitro* modeling for the wider biological research community.

## DEVICE AVAILABILITY

Our goal is to democratize this modular MPS platform for general use. If you’d like to collaborate or have access to the device, please contact us at www.gleghornlab.com/resources.

## ACKNOWLEDGEMENTS

The authors thank Dr. Mark Mirotznik for access to the surface profilometer. Biorender was used in part to create figure 3 and figure 6. This work was supported in part by grants from the National Institutes of Health: U19AI158930 (JPG), T32GM133395 (KMN), and F31HD105398 (KMN).

## DATA AND CODE AVAILABILITY

The data that support the findings of this study are available from the corresponding author upon reasonable request.

## AUTHOR CONTRIBUTIONS

Conceptualization (DJM, KMN, JPG), Methodology (DJM, KMN, FR, BF, JPG), Investigation (DJM, KMN, FR, BF, AMZ, KB), Validation (DJM, KMN, FR, JPG), Visualization (DJM, KMN, FR, BF JPG), Formal Analysis (DJM, KMN, FR, JPG), Writing – Original Draft (DJM, KMN), Writing – Review & Editing (DJM, KMN, FR, BF, AMZ, KB, JPG), Supervision (JPG), Project Administration (JPG), Funding acquisition (KMN, JPG)

